# Exposure to sevoflurane results in changes of transcription factor occupancy in sperm and inheritance of autism

**DOI:** 10.1101/2021.03.08.434461

**Authors:** Hsiao-Lin V. Wang, Samantha Forestier, Victor G. Corces

## Abstract

One in 54 children in the U.S. is diagnosed with Autism Spectrum Disorder (ASD). De novo germline and somatic mutations cannot account for all cases of ASD, suggesting that epigenetic alterations triggered by environmental exposures may be responsible for a subset of ASD cases. Human and animal studies have shown that exposure of the developing brain to general anesthetic (GA) agents can trigger neurodegeneration and neurobehavioral abnormalities but the effects of general anesthetics on the germ line have not been explored in detail. We exposed pregnant mice to sevoflurane during the time of embryonic development when the germ cells undergo epigenetic reprogramming and found that more than 38% of the directly exposed F1 animals exhibit impairments in anxiety and social interactions. Strikingly, 44-47% of the F2 and F3 animals, which were not directly exposed to sevoflurane, show the same behavioral problems. We performed ATAC-seq and identified more than 1,200 differentially accessible sites in the sperm of F1 animals, 69 of which are also present in the sperm of F2 animals. These sites are located in regulatory regions of genes strongly associated with ASD, including *Arid1b, Ntrk2*, and *Stmn2*. These findings suggest that epimutations caused by exposing germ cells to sevoflurane can lead to ASD in the offspring, and this effect can be transmitted through the male germline inter and trans-generationally.

**Summary sentence:** Pregnant mouse F0 females exposed to sevoflurane give rise to F1 males with sociability and anxiety defects. These behaviors are transmitted to F2 and F3 males. Their sperm show changes in transcription factor occupancy in genes implicated in autism.

## Introduction

Autism spectrum disorder (ASD) is a group of neurodevelopmental conditions characterized by deficiencies in social interactions and communication, restricted and repetitive behaviors [1, 2], and a range of comorbid medical conditions including intellectual disability, attention deficits, sensory sensitivities, and gastrointestinal problems. The global prevalence of ASD continues to increase significantly, with more than 1% of individuals affected by ASD and a male to female ratio of 3 to 1 [2, 3]. The complex etiology of ASD includes both genetic and environmental risk factors [1, 2, 4, 5]. Based on twins and multiplex ASD family studies with multiple affected children, the ASD heritability is estimated at ~40 to 90% [2, 6, 7]. Currently there are more than 500 candidate genes recognized by the autism research initiative supported by the Simon Foundation [8] that are significantly associated with clinical diagnosis of ASD. However, these genetic variants can only be found in less than 30% of ASD patients and most of the high risk ASD variants are rare de novo mutations [1]. For example, mutations in the *ADNP* gene, which has been identified as the most commonly found single gene involved in ASD [1] are only present in 0.17% of ASD individuals [9, 10] and only about 205 children diagnosed with ASD have been found to carry mutations in *ADNP* as of January 2019 [11]. In contrast, based on recent larger cohort studies, it has been estimated that environmental risk factors may contribute up to 50% of ASD [5]. Viral infection during pregnancy is the best known environmental risk factor to increase the frequency of autism in the offspring [4, 12, 13]; for example, based on a large Swedish study with a sample size of 24,414 ASD cases, 3.7% of mothers of ASD cases had experienced a viral infection during pregnancy [13]. Therefore, findings in human and animal studies strongly suggests that inherited and de novo mutations may not be the only contributors to ASD pathology, and the contribution of environmental risk factors, possibly through effects on the epigenome, requires careful investigation.

Increasing clinical evidence suggests a significant association between exposure to general anesthetics (GA) alterations in brain development, behavior abnormalities, cognitive impairment, and language deficits [14-16]. Studies in rodents and nonhuman primates have shown that exposure of the developing brain both prenatally and postnatally to commonly used anesthetic agents, including isoflurane and sevoflurane, can result in neurobehavioral abnormalities, including social behavior deficits, increased repetitive behavior and hyperactivity, and memory impairment [14, 16-23]. Many of the deficiencies associated with exposure to anesthetic agents may be long-lasting, leaving irreversible adverse effects. GA-triggered changes in the central nervous result in neuropathologies similar to those observed in ASD patients, including defects in synaptic architecture and abnormal pyramidal neurons of the medial prefrontal cortex [2, 14, 24, 25]. Behavioral phenotypes observed in animal studies also capture impairments in social and repetitive behaviors shown by ASD patients.

It is unclear whether environmental exposures result in epigenetic alterations of the germline that can be transmitted to subsequent generations resulting in ASD [1, 2, 26]. To address this issue, we chose to study to effect of sevoflurane, the most widely used GA agent currently, with a well-established effects on neurotoxicity as a result of direct exposure [14, 16, 19-23]. We exposed pregnant mouse females during the time of development when the embryo is undergoing neurogenesis and the germline undergoes epigenetic reprogramming. Using sociability and repetitive behavior tests, we found that F1 males are significantly impaired in social and repetitive behaviors when compared with unexposed control animals. Furthermore, more than 40% of the F2 and F3 progeny exhibit ASD-like behaviors. We then used ATAC-seq in the sperm of F1 and F2 animals exhibiting ASD behaviors and found 69 differentially accessible sites present in both generations and located in the regulatory regions of ASD-associated genes. These findings suggest that epimutations caused by exposing germ cells to sevoflurane can lead to ASD in the offspring, and these epimutations can be transmitted through the male germline inter and trans-generationally.

## METHODS

### Animals and experimental design

All animal experiments were approved by the Emory University Institutional Animal Care and Use Committee (IACUC). Adult 8-12 weeks old CD-1 IGS mice (Charles River lab, USA) were used for all experiments. Mice were housed in standard cages on a 12 h light/dark cycle and given ad lib access to food and water. Pups were weaned at 21 days old and group housed in sex-matched cages.

Pregnant mouse females were exposed to sevoflurane as outlined in Figure 1A. Healthy 8-12 weeks old CD-1 male and female mice were randomly assigned into two groups, the sevoflurane group and the control group, and timed-matings were conducted. The mating pairs were examined the next morning and the female mice were designated as gestational day 0 (G0) if vaginal plugs were detected. The embryos were designated as in embryonic day 0 (E0.5). The plugged females were separated from the males and individually housed and weighed daily to confirm successful timed matings and pregnancies. Verified pregnant female mice were separated into two groups, control and exposed. Pregnant female mice assigned to the exposed group were exposed to 3% sevoflurane under one of two regiments: (1) seven 2 h exposures daily between E7.5 and E13.5 or (2) single 2 h exposure on E12.5 (Figure 1B). Exposure to sevoflurane was conducted using SomnoSuit (Kent Scientific) with a secure nose cone, starting with a 3 min induction period with 6% sevoflurane at a flowrate of 0.5 L/min supplied in 30% O_2_. Anesthesia was then maintained with 3% sevoflurane at a flowrate of 0.5 L/min for 2 h. Exposed pregnant females were allowed to recover in 30% O_2_ for 15 min in the chamber and monitored for 30 min with access to moist food. Pregnant female mice assigned to the control group were never exposed to sevoflurane and were bred in the same animal facility. All exposed and control dams were allowed to give birth naturally and the postnatal body weights of pups were measured bi-weekly from postnatal weeks 6 to 12.

**Figure 1.**
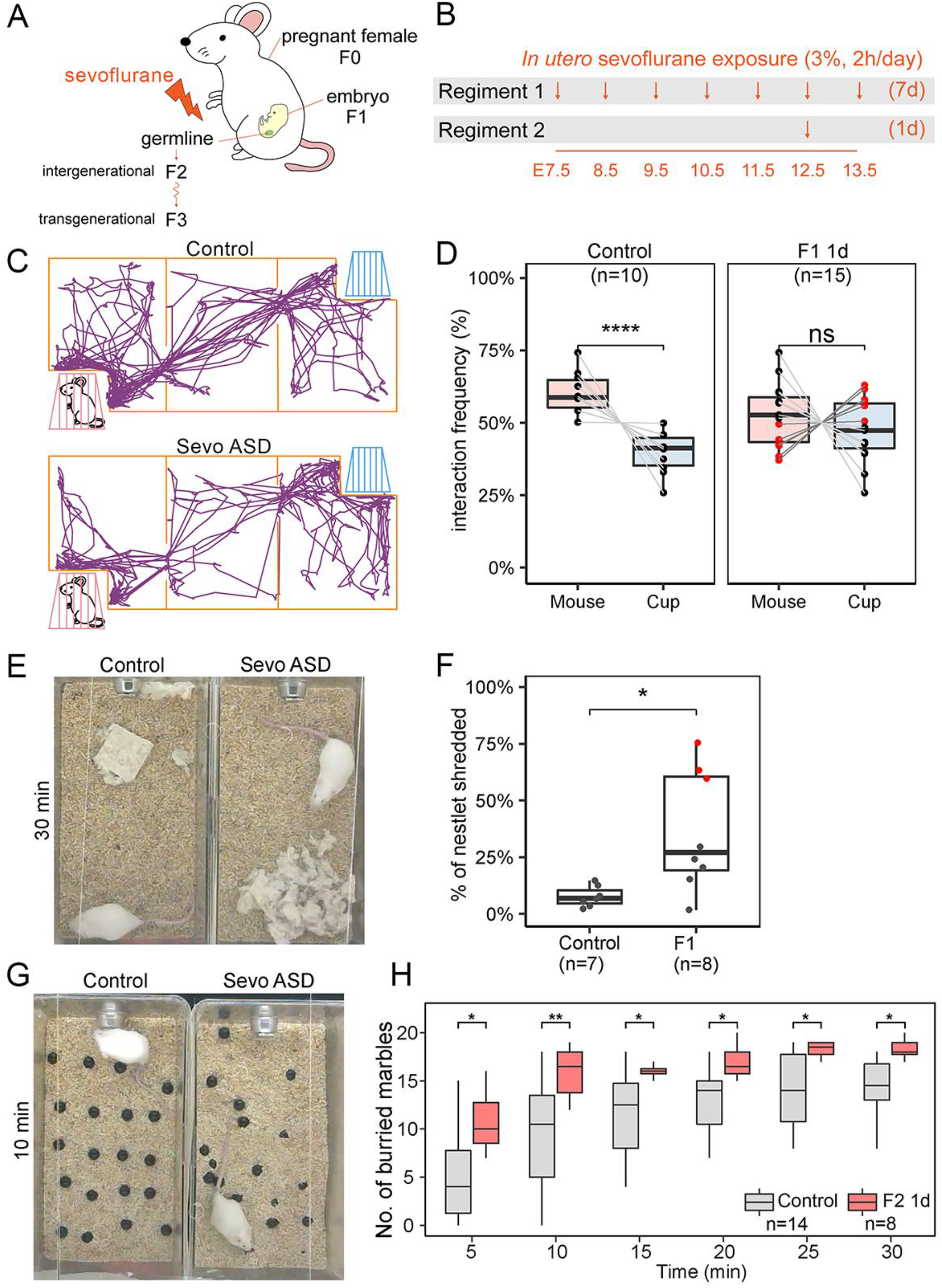
*In utero* exposure to sevoflurane during brain development results in ASD-like phenotypes. (A) Diagram of sevoflurane exposure *in utero* and possible effects on subsequent generations. (B) Diagram of the two exposure regiments utilized in the experiments. One regiment involves daily 2 h exposures for seven days starting at E7.5 and ending at E13.5. A second regiment involves a single 2 h exposure at E12.5. (C) Schematic representation of the three-chamber sociability test with a typical example of the movements of a control and a sevoflurane-exposed ASD mouse (Sevo ASD). (D) Results from the sociability test in F1 male mice exposed to sevoflurane using the 1-day regiment. Control male mice (n=10) prefer the chamber housing an unfamiliar stranger mouse (one-way ANOVA, p< 0.0001) whereas F1 mice (n=15) show no statistically significant differences in their preference for the empty cup or the unfamiliar mouse. Red dots represent mice with sociability index (SI) <0 and ASD phenotypes whereas black dots represent mice with sociability index (SI) >0 and normal phenotypes. (E) Photographs of control and sevoflurane ASD animals at the end of the nestlet shredding test, used to quantify the degree of repetitive and compulsive behaviors, showing differences in shedding behavior between the two. (F) Percentage of shredded nestlet (y-axis) showing the distribution of nestlet shredding for control (n=7), and F1 male mice exposed using the 1-day regiment. Extreme outliers are marked in red, indicating individuals with ASD-like behavior (one-way ANOVA, *p < 0.05). (G) Pictures of control and sevoflurane ASD animals after 10 min of the marble burying test showing the number of marbles buried by each mouse. (H) Results from the marble burying tests conducted with control and F1 mice exposed to sevoflurane using the 1-day regiment. The x-axis indicate time passed since the beginning of the test (one-way ANOVA; **p < 0.01, *p < 0.05).

The progeny from the sevoflurane-exposed females were crossed as outlined in Figure S1A and at least two independent biological replicates were performed per condition. Exposed pregnant females were designated as F0, and the embryos exposed *in utero* were referred to as F1. F1 sevoflurane males were bred to unexposed control CD-1 females to obtain the F2 paternal outcross generation. F2 males were crossed to unexposed control CD-1 females to obtain the F3 paternal outcross generation. Unexposed CD-1 mice were used as control groups and control animals were bred in the same manner as the sevoflurane cohort for each generation. No sibling breeding was performed.

### Behavioral tests

All male animals were evaluated for their behavioral phenotypes after they reached sexual maturity at the age of 7 to 15 weeks following well established protocols [27-29]. Only males were used for ASD-like behavioral tests based on published recommendations [30, 31], since female mice show higher variability in behavioral tests perhaps due to hormonal fluctuations during the estrus cycle [30]. For all tests, 1) age-matched control animals were utilized; 2) test animals had a minimum 7-day break between each behavioral test; 3) test mice were placed in a new cage 24 h before testing; 4) all tests were conducted between 9 am and 4 pm during the light phase of the light-dark cycle; 5) mice were acclimated in a designated animal behavior room without disturbance for at least 60 min before tests; 6) all materials were disinfected between each trial; 7) food and water were withheld during testing. The order of tests was as follows when possible: three-chamber sociability test, nestlet shredding, and marble-burying test. All test results were analyzed by a blinded observer lacking information on the identity of the animals to ensure absence of possible biases.

#### Three-chamber sociability test

The three-chamber sociability tests were performed as described previously [32]. The apparatus consists of three equal sized chambers 40 x 40 cm each separated by clear acrylic walls with doors cut out (Figure 1C). Both side chambers contain a wired cup in opposite corners with a 1 L glass bottle on top to prevent the testing mouse from climbing. The test consists of two 10 min sessions. In the first session, the mouse being tested is able to habituate to the apparatus by freely exploring all 3 chambers with empty cups in both side chambers. In the second session, the test mouse is gently confined to the center chamber and a wild-type age and sex matched unfamiliar stranger mouse is placed in one of the two wired cups; the test mouse is then allowed to freely explore all three chambers for 10 min. The movements of the test mice were recorded using a digital video camera and the distance traveled and time spent in each chamber were analyzed using ANY-MAZE (Stoelting Co., Wood Dale, IL, USA) (Figure 1C). To interpret the results of the sociability test, the time each mouse spent exploring the chamber with the stranger mouse or the chamber with the empty cup were quantified. A sociability index (SI) was used to compare the significance of differences between ASD and control animals [33]. Animals were classified as ASD when SI < 0, whereas mice showing SI > 0 were defined as non-ASD.

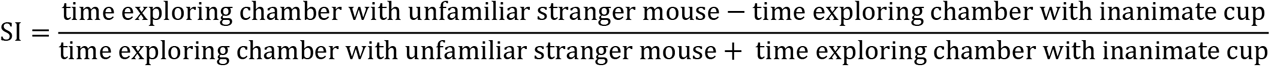

#### Nestlet shredding test

The nestlet shredding tests were performed as described previously [34]. The test mice were housed in a new standard cage for 24 h prior to the test with normal bedding but no nestlet materials. Subsequently, the test mice were placed individually in another clean cage (18 x 32 x 14 cm) containing a pre-weighed unscented square cotton nestlet (start weight). Commercially available cotton nestlets (5 cm × 5 cm, 5 mm thick, ~2.5 g each) were weighed on an analytical balance. The mice were left undisturbed for 30 min after which debris and cage bedding were removed from the portion of un-shredded nestlets. Each nestlet was then weighed (end weight) and scored as described previously [34]. The fraction of shredded nestlet was obtained by dividing the end weight by the start weight and subtracting from one.

#### Marble burying test

Test was performed as previously described [34, 35] in a standard cage (18 x 32 x 14 cm) filled with approximately 4.5 cm of fresh bedding (Bed-o’Cobs, 1/8”). Twenty black marbles 15 mm in diameter were arranged equidistantly on the surface of the bedding in a 4 x 5 arrangement. The test mice were placed gently in the corner of the cage to avoid displacing any marbles and tested for 30 min undisturbed. The movements of test mice were recorded on video during the entire duration of the procedure. The number of marbles buried at different time intervals were counted by blinded observers by analyzing the recorded videos. The marbles were considered buried if >2/3 of the marble surface was covered by bedding material.

#### Data analyses

All animal behavioral tests were scored by statistical analyses and were conducted on raw data using ggplot2 [36] in R version 3.6.3 [37] with two-tailed t-test, one-way ANOVA, or two-way repeated-measures ANOVA. P-values <0.05 were regarded as statistically significant (ns: p > 0.05, *p ≤ 0.05, **p ≤ 0.01, ***p ≤ 0.001, ****p ≤ 0.0001). Box plots were used to report the distribution of behavioral test results and the median and the interquartile range (IQR) between the lower quartile (Q1, 25%) and the upper quartile (Q3, 75%) are shown in the figures. Data points that fall above or below the Q1 - 1.5 * IQR or Q3 + 1.5 * IQR were considered outliers.

### Isolation of sperm

Euthanasia was performed by cervical dislocation and the epididymis was removed. Mature sperm were collected from the dissected cauda epididymis of 12 to 16 week-old adult male mice as previously described [38-41]. Briefly, blood vessel and adipose tissue were removed by dissection, and the cauda epididymis was rinsed with PBS and placed in a small cell culture dish containing freshly prepared Donners medium. The cauda epididymis was punctured with a needle before transferring to a round-bottom tube and incubated at 37°C for 1 h to allow the sperm to swim up. Purity of sperm was examined under a light microscope and was determined to be at least 99.9% if less than one somatic cell was detected after counting 1000 sperm. Numbers of pure sperm collected were determined using a hemocytometer.

### Assay for ATAC-seq

Omni-ATAC-seq was performed as previously described [40-42]. Briefly, a total of 100,000 sperm were used per biological replicate. The nuclei were isolated in digitonin containing buffer (10 mM Tris-HCl pH 7.4, 10 mM NaCl, 3 mM MgCl_2_, 0.1% NP40, 0.1% Tween-20, and 0.01% digitonin) by brief incubation on ice for 3 min. Sperm nuclei were pelleted and resuspended in transposase reaction mix containing 0.05% digitonin and incubated for 30 min at 37°C. After the transposase reaction, the transposed samples were treated with Proteinase K in 0.1 M Tris-HCl, 0.2 M NaCl, 5 mM EDTA, 0.4% SDS, and 0.4 μg/μL Proteinase K at 55°C for 2 h. The genomic DNA was isolated by phenol:chloroform:isoamyl alcohol extraction and ethanol precipitation. ATAC-seq library amplification was done using the KAPA SYBR FAST qPCR Master Mix (2X) Kit (Kapa Biosystems) and 1.25 μM indexed primers using the following PCR conditions: 72°C for 5 min; 98°C for 30 s; and 11 cycles at 98°C for 10 s, 63°C for 30 s, and 72°C for 1 min.

### ATAC-seq data processing

All libraries were sequenced using an Illumina NovaSeq 6000 instrument in a 50 bp paired-end format. Paired reads were aligned to the mouse mm10 reference genome using Bowtie2 [43] with default parameters, except -X 2000. PCR duplicates were removed using Picard Tools [44]. To adjust for fragment size, we aligned all reads as + strands offset by +4 bp and – strands offset by −5 bp [45]. Reads were separated by size into 50-115 bp fragments, which correspond to regions bound by transcription factors (TFs), and 180-247 bp fragments, which correspond to mono-nucleosomes [40, 41, 46]. We will refer to ATAC-seq peaks corresponding to the presence of TFs as THSSs (Tn5 hypersensitive sites). Peaks were called in the THSS and mono-nucleosome fractions using MACS2 with default parameters [47]. ATAC-seq peaks that overlap with the ENCODE blacklist for mm10 were excluded from further analysis [48].

Differential THSSs were analyzed using DiffBind, and regions ±75 bp from the summits were compared [49, 50]. THSS bedgraphs/bigwig files were made using the deeptools bamCoverage function with default parameters except --binSize 1 --normalizeUsing RPGC --ignoreForNormalization chrX chrY chrM -p 25 [51]. Heatmaps of identified differential THSSs were analyzed using the following method. Summits of each THSS were used as anchors and reads mapped to each THSSs ±2 kb from its summit were quantified and binned. To normalize the datasets, the raw read counts were divided by library sizes in millions to obtain reads per million per covered bin (RPMPCG or RPM), which was then visualized with EnrichedHeatmap version 1.15.0 [52]. K-means clustering was done using ComplexHeatmap version 2.3.2 [52].

### TF motif occurrences and TF motif cluster enrichment analysis

Position weight matrix-based TF motif mapping analysis was conducted using MOODS [53]. TF motif cluster, termed TF archetype, analysis was adapted from [54] using the published motif clustering annotation matrix for the mm10 genome. Monte Carlo permutation tests for randomness [55], where the p-value estimates the significance of TF archetype enrichment genome-wide, were conducted over 1000 randomly simulated trials for each TF archetype. The p-value was quantified by the number of randomized trials that are found greater than observed, and p-value ≤ 0.001 were considered statistically significant. As an independent test, the randomization was also tested against the list of 339,815 candidate cis-regulatory elements (cCREs) mapped to the mouse mm10 genome. This list is curated and annotated by ENCODE based on more than 170 DNase-seq and CTCF, H3K4me3, and H3K27ac ChIP-seq datasets from mouse cell lines and primary tissues [56].

### Gene and disease ontology and ASD gene enrichment analyses

Disease ontology enrichment analysis was performed in the Comparative Toxicogenomics Database (CTD) using default parameters [57]. The significance of enrichment is calculated by the hypergeometric distribution and adjusted for multiple testing using the Bonferroni method for each enriched term. The gene ontology for cellular components was performed using PANTHER GO-SLIM [58]. The terms were considered significant if p-value < 0.001. Monte Carlo permutation tests for randomness [55], where the p-value estimates the significance of ASD genes found in the set of differential THSSs, were conducted over 1000 randomly simulated trials for each dataset. P-values ≤ 0.01 were considered statistically significant. The gene ontology enrichment network analysis was performed using GOnet [59] and Cytoscape [60] and p-value < 0.01 were considered statistically significant. Each node of the network consists of genes identified in/around THSSs and is linked by an edge to associated enriched GO terms. The mouse cerebral cortex RNA expression data in TPMs was used to color each node using Mouse ENCODE project dataset and color genes by expression function on GOnet [59, 61, 62].

### Data availability

The raw sequence for ATAC-seq data reported in this paper is deposited in NCBI’s Gene Expression Omnibus (GEO) under accession number GSE167500. Reviewers can access these data using token ifyfkioobhkjzwf. Custom scripts used to separate ATAC-seq reads into subnucleosomal (THSSs) and nucleosome-size ranges and the other custom scripts used in data analyses are available upon request without restrictions.

## Results

### Prenatal exposure to sevoflurane during embryonic neurogenesis results in ASD-like behaviors

Previous studies have reported a lasting impact of GA on the human brain and in animal models, including a high frequency of inheritance of these effects [14-24, 26]. Little is known about the mechanisms by which GA exposure results in adverse effects considering how commonly patients of different ages and health status are exposed to GA. Therefore, identifying the effects of the duration and timing of GA exposure may be important to understand the nature and extent of GA induced phenotypes. Currently, two exposure regiments have been mainly used to study the effect of GA by exposing either pre-natal or post-natal animals to clinically relevant conditions. Several pioneer studies have focused on exposing rodent neonates and animals less than 17 days old undergoing rapid synaptogenesis [18-20, 24]; while only recently a few studies have begun dissecting the effect of GA on the fetus during different periods of gestation in pregnant females [21-23]. To understand the impact of general anesthetics on developing embryos exposed *in utero*, specifically the effect on the germline and on neural development, we exposed pregnant CD-1 females to clinically relevant dosage and duration of sevoflurane, either daily during E7.5-E13.5 or once during E12.5 (Figure 1 A,B). We will refer to these two different exposure regiments as 7-day and 1-day exposures. The rationale for the timing of the two exposures is as follows. The epigenome of primordial germ cells (PGCs) is reprogrammed, including erasure of most DNA methylation, starting at E7.5 and ending at E13.5. This time may represent a window of special vulnerability to epigenetic changes caused by exposures to chemicals in the environment. In addition, the different cell types of the brain differentiate during this time and, for example, genetic perturbation in the expression of genes involved in autism during E12.5 results in ASD phenotypes in mice [63]. In support of this, exposure of pregnant females to BPA during the window of germline reprogramming and differentiation leads to heritable obesity phenotypes in the unexposed F3-F5 generations [39, 64-66].

F1 animals exposed *in utero* to sevoflurane are born without overt morphological phenotypes. Females from the control and sevoflurane exposed groups have the same 21-day gestation and deliver similar number of 8-15 pups with no significant variation in sex ratio. The behavior of control and mice exposed *in utero* with the 1-day regiment was assessed at 7-10 weeks of age. Control male animals exhibit consistent behavioral phenotypes whereas control females showed significant variability in social behavioral tests. Therefore, we only used males to assess the possible effects of sevoflurane on behavior. We used the three-chamber test to assess sociability [4, 32] and only examined F1 males subjected to the 1-day exposure regiment (Figure 1D). Similar to other studies in which embryos were exposed *in utero* to GA [21-23], we found that F1 7-10 weeks old males exhibit impairment in social and repetitive behaviors when compared to unexposed males. We first evaluated all F1 1-day exposed males and found that, based on the medium interaction frequency, there is no significant differences in the preference of these mice for another unfamiliar stranger mouse or the empty cup (p-values >0.05, Figure 1D), suggesting that a single 2 h sevoflurane exposure at E12.5 is sufficient to cause social interaction deficits. To further dissect the information resulting from these sociability measurements, we separated the F1 exposed mice into two groups. Males showing a higher preference for the empty cup (sociability index, SI, < 0; −1 to −26, average - 15) were classified as ASD mice and represented in the figures by red dots. Mice showing a preference for the unfamiliar stranger mouse (SI > 0; 3 to 35, average 16) were classified as non-ASD mice and represented in the figures as black dots. Based on this classification, 40% (n= 6 out of 15) of F1 males exhibit stronger preference towards an empty cup than to an unfamiliar stranger mouse and are classified as ASD (Figure 1D). We then used the nestlet shredding test to assay for repetitive and compulsive-like behaviors [34], another key impairment commonly identified in ASD patients. Figure 1E shows an example of the behavior in this test of a control mouse and a moused classified as ASD based on the sociability test shown in Figure 1D. We found that adult F1 males show a statistically significant increase (p-value < 0.05) in nestlet shredding compared to age and sex-matched control animals (Figure 1F). Among all exposed F1 males tested, 37.5% (n=3 out 8) exhibit an extreme increase in nestlet shredding (red dots, nestlet shredding score above the upper quartile, in Figure 1F). Two of three animals with extreme shredding phenotypes are also impaired in the sociability test shown as red dots in Figure 1D. To confirm these results, we performed the marble burying test, which also measures repetitive and compulsive behaviors (Figure 1G). We found that, independent of the test duration, F1 males exposed to sevoflurane *in utero* using the 1-day regiment buried significantly more marbles than unexposed controls (Figure 1H). Therefore, results from three independent behavioral tests suggest that 37-40% of the F1 male progeny of sevoflurane-exposed F0 pregnant females exhibit ASD-like phenotypes with both social interaction deficits and increased repetitive behaviors.

### ASD phenotypes caused by prenatal exposure to sevoflurane during germ cell reprogramming can be transmitted transgenerationally

In addition to affecting brain development, sevoflurane exposure using either the 1-day or 7-day regiments may also affect the gametes, since the time window used for the exposure corresponds to the time when the epigenetic content of the germline is reprogrammed. To examine whether exposure to sevoflurane causes epigenetic alterations in the gametes that can be transmitted to subsequent generations and elicit ASD-like phenotypes, we crossed F1 males exposed to sevoflurane for 2 h daily using 1-day and 7-day regiments with unexposed control females and assessed the behavior of the F2 adult male offspring. To analyze the heritability of sevoflurane induced sociability impairments specifically, for crosses involving F1 males exposed for 1 day, we were able to separately cross ASD (red dots in Figure 1D) and non-ASD (black dots in Figure 1D) males to control females. We will refer to the F2 progeny of the former cross as F2-ASDp to reflect the fact that the phenotype was transmitted paternally from an ASD male, and to the F2 progeny from the latter cross as F2-nASDp to indicate that the F1 father was non ASD. F2-ASDp mice show no significant difference in the preference for the stranger mouse or empty cup, with 47% (8 out of 17) showing SI < 0 and an ASD phenotype, suggesting that the germline of their F1 fathers was epigenetically altered by sevoflurane exposure, and that these alterations can be transmitted to the somatic cells of the embryo to elicit ASD symptoms (Figure 2A). In contrast, F2-nASDp males show a preference for the unfamiliar stranger mouse over the empty cup, suggesting that these mice do not have sociability defects and that the germline of their F1 non-ASD fathers was not affected by exposure to sevoflurane (Figure 2A). Results from the nestlet shredding test (Figures 2B,C) and from the marble burying test (Figure 2D) confirm these conclusions.

**Figure 2.**
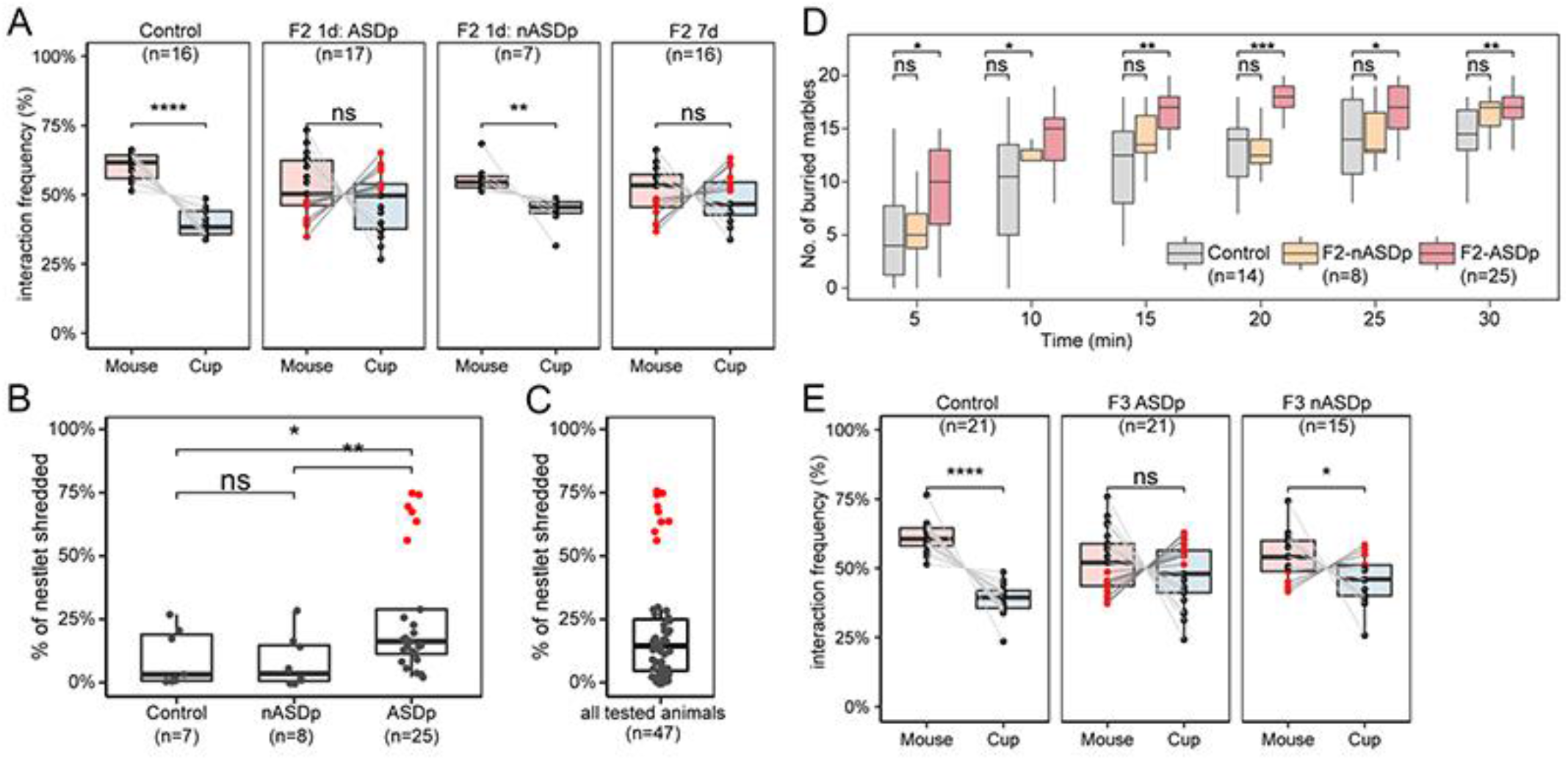
Exposure to sevoflurane *in utero* during germline reprogramming results in paternal inter- and trans-generational transmission of ASD phenotypes. (A) Results from the sociability test in F2 male progeny arising from F1 fathers exposed to sevoflurane while *in utero* using the 1-day exposure regiment and either showing ASD phenotypes (ASDp) or showing normal behavior (nASDp). The rightmost panel shows results for the F2 mice progeny of F1 males subjected to the 7-day exposure regiment while *in utero*. Red dots represent mice with sociability index SI<0 and ASD phenotypes whereas black dots represent mice with SI>0 and normal phenotypes. (B) Percentage of shredded nestlets (y-axis) to quantify the degree of repetitive and compulsive behaviors. Nestlet shredding data of control, F2 sired by 1-day exposed F1 with no ASD phenotypes (nASDp) and F2 sired by 1-day exposed F1 with ASD phenotypes (ASDp). F2 ASDp males show a significant increase in nestlet shredding. A group of animals classified as ASD are marked by red dots. (C) Distribution of nestlet shredding for all tested animals (n=47). Mice are clearly separated into two groups and the extreme outliers marked in red are classified as ASD individuals. (D) Marble burying tests were conducted in control, F2 sired by 1-day exposed F1 males with normal behavior (nASDp) or ASD behavior (ASDp). (E) Results from the sociability test in F3 male progeny arising from F2 fathers whose F1 fathers were exposed to sevoflurane while *in utero* using the 7-day exposure regiment and either showing ASD phenotypes (ASDp) or showing normal behavior (nASDp). Red and black dots are as in previous panels. For all panels, statistical tests were performed using one-way ANOVA (****p < 0.0001, ***p < 0.001, **p < 0.01, *p < 0.05, ns, not significant).

The behavioral phenotypes of F1 males exposed for 7 days was not assessed and, therefore, the ASD status of males involved in these crosses was not known. These F2 mice (F2 7d) show on average no preference for the stranger mouse or the empty cup, and 44% (7 out of 16) of them show a SI < 0 and an ASD phenotype (Figure 2A). Based on this result, ASD (red dots) and non-ASD (black dots) F2 mice from the 7-day exposure regiment were crossed with control unexposed females and the F3 progeny was examined using the sociability test. Females from both crosses delivered similar numbers of F3 pups with consistent 21-day gestation periods and no statistically significant variation in the sex ratio of the progeny. We found that adult male F3 animals sired by ASD F2 males exhibit impairment in sociability (Figure 2E).

More than 48% of these social interaction impaired F3 males exhibit the same preference towards an inanimate object than for an unfamiliar age and sex matched stranger mice (SI < 0; n= 10 out of 21). In contrast, adult F3 males sired by non-ASD F2 animals exhibit normal sociability behavior (p-value < 0.05), and 73% of all tested animals preferentially interact with an unfamiliar age and sex matched mouse instead of an inanimate object (SI > 0; n=11 out of 15) (Figure 2E). Therefore, these observations suggest that *in_-utero* exposure of mouse embryos to human relevant doses of sevoflurane can lead to inter- and trans-generational transmission of ASD-like phenotypes, suggesting that exposure to sevoflurane alters the epigenetic content of the male germline.

### *In utero* exposure to sevoflurane results in changes in transcription factor distribution in F1 sperm

Mammalian sperm contain chromatin in which most histones have been replaced for protamines. However, around 8% of the histones found in somatic cells remain in mature sperm and these histones carry a variety of active and inactive histone modifications [40]. Furthermore, sperm DNA is methylated, and many other chromatin proteins, including RNA polymerase II and other components of the transcription complex present at promoters and transcription factors (TFs) bound to enhancers can also be found in mammalian sperm [41]. Since exposure to sevoflurane results in ASD-like phenotypes that can be paternally transmitted to the unexposed F2 and F3 offspring, we reasoned that the sperm of ASD F1 mice likely carry epimutations responsible for this transmission. Based on previous results, suggesting that patterns of TF occupancy in sperm chromatin are maintained in the pre-implantation embryo whereas histone modifications and DNA methylation are at least partially erased after fertilization [41,67], we hypothesized that exposure to sevoflurane may alter the distribution of TFs at specific regions of the sperm genome. To test this hypothesis, we performed ATAC-seq in sperm isolated from the cauda epididymis of control and sevoflurane 1-day and 7-day exposed F1 male animals [41, 42]. We separated the reads of each ATAC-seq experiment and focused specifically on sub-nucleosome size reads less than 115 bp in length, which correspond to DNA protected from Tn5 insertion by bound TFs (THSSs) at enhancers and promoters. We performed two independent pair-wise comparisons of THSSs to identify differences in TF binding using stringent cut-off criteria, FDR <0.05, 2-fold change, and >4 counts per million per condition.

We examined differences between 1) F1 sevoflurane 1-day exposure and control sperm and 2) F1 sevoflurane 7-day exposure and control sperm. We found that the two samples from exposed sperm correlate with each other better than with control (Figure 3A). Furthermore, we find 1,019 and 1,010 differential THSSs in the two comparisons, respectively (Figure S2), suggesting that the occupancy of a large number of TFs was altered as a result of exposure of PGCs to sevoflurane.

**Figure 3.**
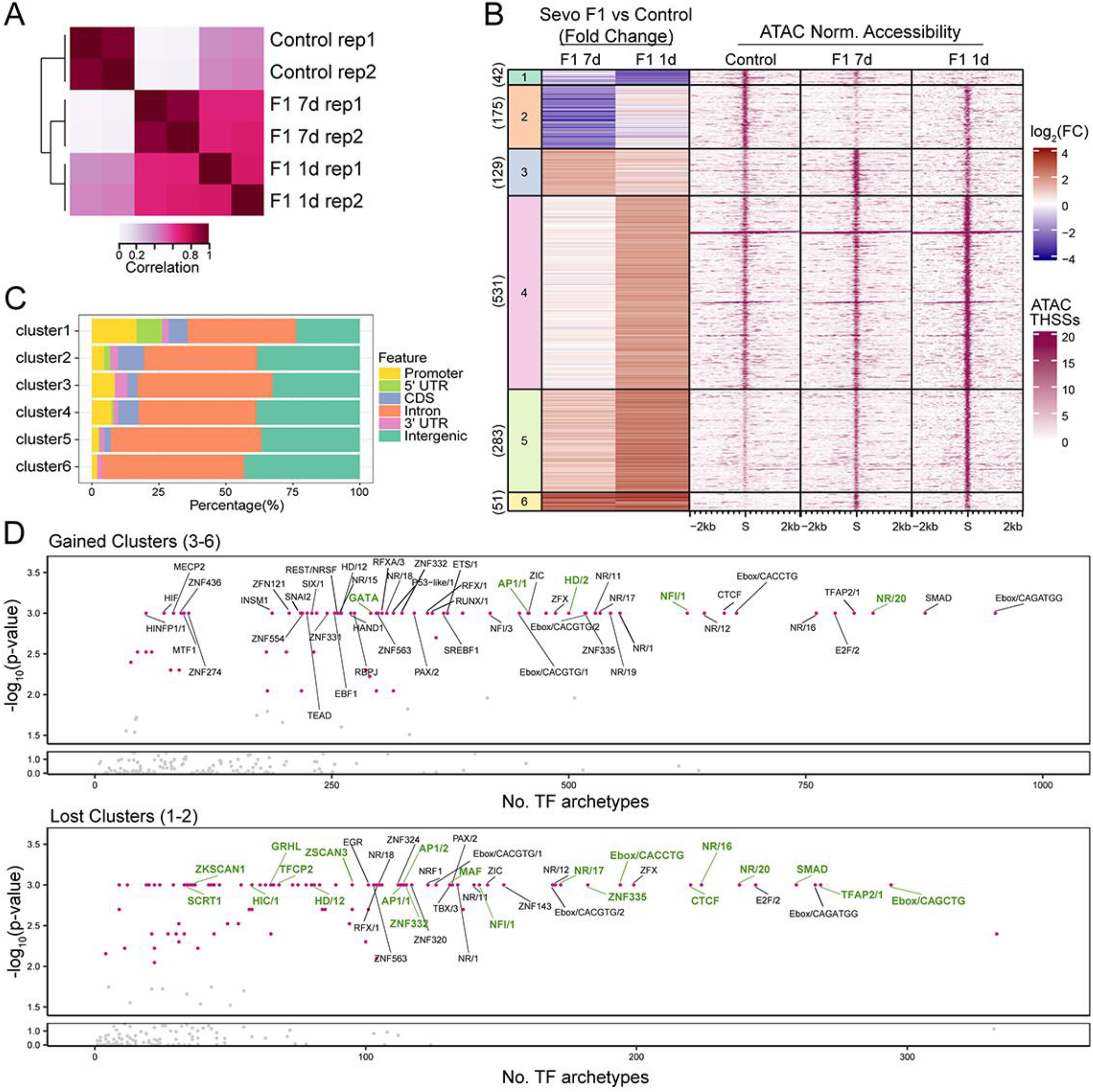
Sevoflurane induced alterations in TF distribution in sperm of F1 animals. (A) Pearson correlation heatmaps of read coverage at differential THSSs when comparing sperm ATAC-seq from control, 7-day exposed, and 1-day exposed F1 animals. THSSs are defined as regions ±75 bp from the summits. (B) k-means clustering analysis of differential THSSs (n=1211) using subnucleosome-size reads from ATAC-seq experiments. Fold change of pair-wise comparisons (shown on the left) of 7-day exposed F1 vs control and 1-day exposed F1 vs control were used in k-means clustering. The corresponding normalized subnucleosomal reads are shown on the right for control, 7-day exposed F1, and 1-day exposed F1 datasets. (C) Genome-wide distribution of THSSs present in the different clusters shown in panel B; promoters were defined as TSS-0.5 kb. (D) TF motif archetype enrichment analysis at differentially accessible clusters from panel B showing increased accessibility (clusters 3, 4, 5, and 6; top) and lost accessibility (clusters 1 and 2; bottom). TF motif archetype clusters with p-value less than 0.001 based on Monte Carlo permutation tests simulated genome-wide were labeled with a pink dot. TF motif archetypes with p-value < 0.001 in the permutation tests simulated within the pool of ENCODE’s candidate cis-regulatory elements (cCREs) were marked in green.

Since F1 males from both 7-day and 1-day exposure regiments give rise to F2 progeny with ASD phenotypes, we combined the differential THSSs identified in each comparison, performed k-means clustering, and identified six separate clusters of differential THSSs (Figure 3B). Clusters 1 and 2 show a decrease in Tn5 accessibility (loss of TF binding) whereas clusters 3-6 exhibit gain of Tn5 accessibility (gained TF binding) when comparing sperm from exposed F1 versus control animals. In the case of cluster 4 this only applies to F1 males exposed to the 1-day regiment. These differential THSSs are found in introns and intergenic regions, suggesting that they correspond to regulatory sequences (Figure 3C). To gain insights into the nature of the TFs presents at differential THSSs, we performed position weight matrix-based TF motif mapping [54]. To reduce the redundancy of TF motif models, we adapted a recently published strategy to group individual TFs with similar sequence motifs into clusters termed TF archetypes [54]. We then analyzed the total number of occurrences for each TF archetype instead of specific TFs. We also subjected each TF archetype to Monte Carlo permutation tests for randomness [68], where the p-value estimates the significance of TF archetype enrichment (Figure 3D). Interestingly, the CTCF, Ebox, and nuclear receptor archetypes were among the most abundant found in both the gained and lost clusters (p-value <0.001). Specifically, NR/20 archetypes, which consist of androgen receptor (Ar) and glucocorticoid receptor (GR) binding motifs, is highly enriched in differential THSSs clusters (Figure 3D). The NR/20 archetype is found in 821 gained and 238 lost clusters (Figure 3D). Ar interacts with DNA when it is bound by androgens such as testosterone [69, 70], and it has also been shown that testosterone levels are positively correlated with Ar gene expression [71]. Supplements of testosterone increase Ar gene expression in human studies [72, 73]. These observations are interesting in the context of published evidence showing that exposure to sevoflurane increases testosterone levels in male animals [74, 75] and that a surge of testosterone takes place during germ cell and gonadal sex differentiation [70]. It is therefore possible that Ar and related nuclear hormone receptors are responsible for bookmarking regions responsible for sevoflurane induced epimutations.

### Sevoflurane induced alterations of TF binding in F1 sperm are enriched in genes involved in nervous system disease and mental disorders, many of which are strong ASD candidate genes

The genomic location of differential THSSs could shed light on the genes and pathways involved in sevoflurane induced ASD pathologies. To address this question, we examined the nature of the genes located at or near differential THSSs identified in the sperm of F1 sevoflurane-exposed with respect to control animals. We performed disease ontology enrichment analysis (hypergeometric distribution, p-value < 0.01) and found that diseases of the nervous system and mental disorders were among the most enriched terms found in both gained and lost differential THSSs clusters (Figure 4A). We also performed enrichment analysis by comparing the proximity of differential THSSs to strong ASD candidate genes using the list of 533 human genes with mouse homologs in the SFARI category S, 1 and 2 [8]. We found that both gained and lost differential THSSs are enriched at these ASD candidate genes (p-value = 0.001 and 0.025 for gained and lost clusters, respectively), including *Arid1b, Ntrk2*, and *Sox6* (Figure 4B-D and Figure S3). Interestingly, the differential THSSs located at these ASD candidate genes harbor motifs for NR/20 and/or CTCF TF archetypes (Figure 4B-D) that match to the consensus motifs model of Ar and Ctcf (Figure 4E). Eighteen of these ASD genes are expressed in adult mouse cerebral cortex with greater than 5 TPMs (transcripts per million) and involved in similar biological processes, including nervous system development, neurogenesis, and chromatin organizations (Figure S3). Therefore, it is likely that these differential THSSs represent functional regulatory elements of genes that play a role in causing ASD phenotypes.

**Figure 4.**
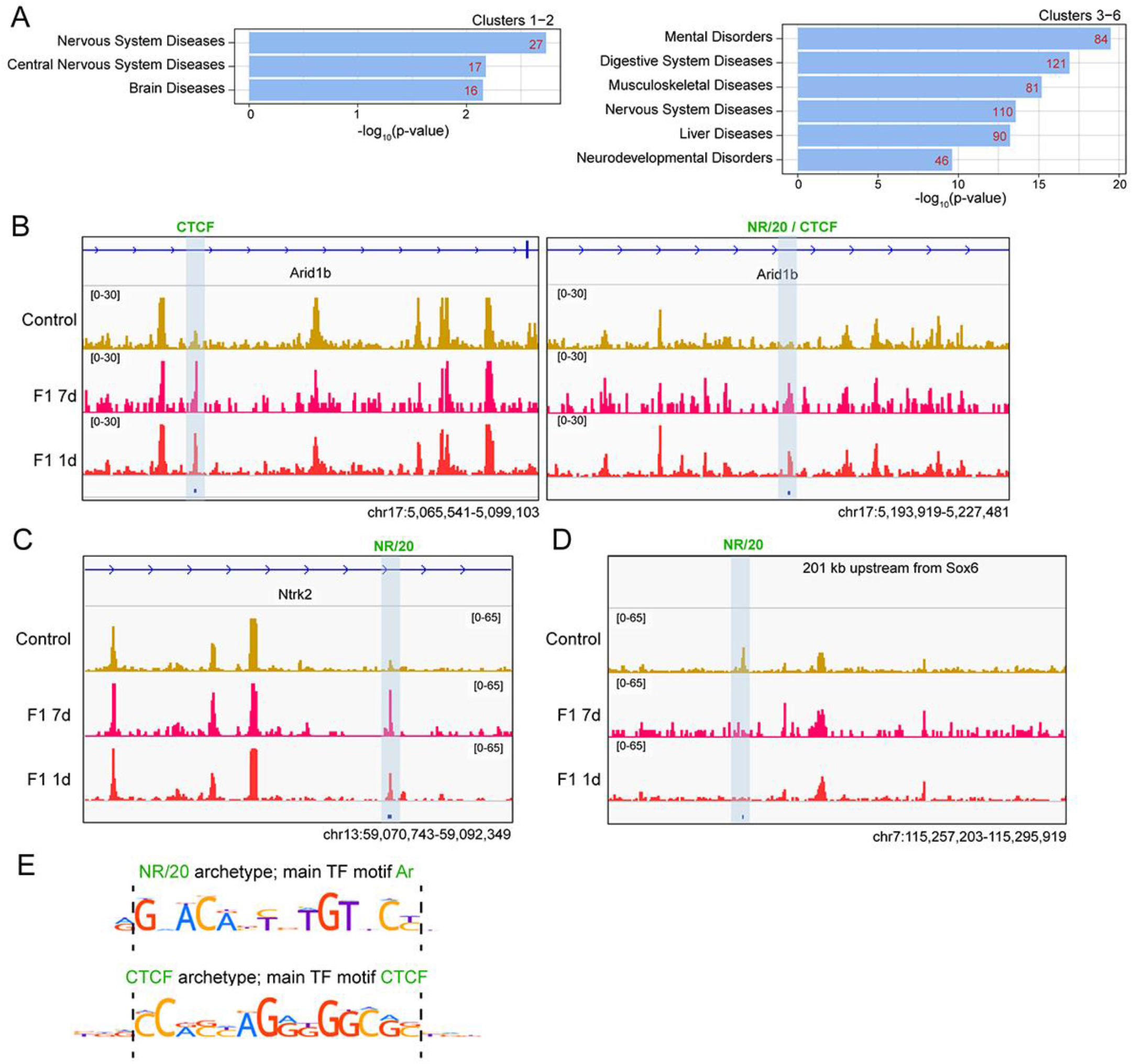
Sevoflurane induced alterations of TF binding in F1 sperm are enriched in or around genes involved in nervous system diseases and mental disorders. (A) Genes located on or closest to differential THSSs from the different clusters shown in Figure 3B were subjected to disease ontology enrichment analysis using the Comparative Toxicogenomics Database (p-value <0.01, hypergeometric distribution with Bonferroni correction). (B-D) Genome browser views of differential THSSs (highlighted by blue shaded box) with NR/20 and/or CTCF TF archetypes in/around ASD genes: (B) *Arid1b*; (C) *Ntrk2*; and (D) *Sox6*. Normalized counts per million (CPM) are shown for control, F1 7-day, and F1 1-day datasets. These sites contain binding motifs for Ar and CTCF. (E) Position weight matrices for Ar and CTCF present at differential THSSs shown in panels B-D.

### Sevoflurane induced ASD-like phenotypes can be inherited trans-generationally through the male gametes

Based on the animal behavior data and design of the crosses, we hypothesize that sperm from F2 ASD males is solely responsible for transmitting the ASD-like phenotypes, since the sperm from these mice were never exposed to sevoflurane (Figure 1A and Figure S1), yet it was sufficient to sire F3 animals with strong ASD-like phenotypes when crossed with control females. Based on our findings above analyzing the alterations of TF binding in the sperm of F1 ASD males, we reasoned that differential THSSs found in F1 ASD sperm should remain in the sperm of F2 ASD animals, since their F3 progeny shows ASD phenotypes. To address this possibility, we performed ATAC-seq using sperm isolated from the F2 ASD males from the 7-day exposure regiment and analyzed sub-nucleosome size reads that represent sites bound by TFs. We identified 1,281 differential THSSs when comparing sperm from F2 ASD and control animals with stringent cut-off criteria (FDR <0.05, >2 fold change, and >4 counts per million per conditions), with significantly more gained than lost THSSs, even though the germline of F2 ASD animals was never exposed to sevoflurane (Figure S4A).

To identify altered TF sites responsible for the trans-generational inheritance of sevoflurane-induced ASD-like phenotypes, we compared the differential THSSs found in the sperm of F1 and F2 ASD animals from the 7-day exposure regiment. We identified 69 differential THSS sites that are present in both the sperm of F1 and F2 ASD but not in control mice and performed k-means clustering of these sites into 5 clusters (Figure 5A). Clusters a and b exhibit loss of TF binding in both F1 and F2 ASD animals with respect to control. In contrast, cluster c, d, and e exhibit gain of TF binding. These 69 sites are located in introns and intergenic regions (Figure 5B) and enriched in introns and intergenic regions (Figure 5C). These putative regulatory sequences are enriched near genes involved in mental disorders, nervous system diseases and ASD (p-value < 0.01) (Figures 5D and S4B). Interestingly, some of the genes adjacent to THSSs with increased occupancy are expressed in the cerebral cortex (Figure S4B) and encode proteins located at distal axons, cytoplasmic and intracellular vesicles, and synapses (GO analysis - cellular components, p-value < 0.01) (Figure 5E). These observations suggest that these 69 sites may represent functional regulatory elements responsible for the transgenerational transmission of sevoflurane-induced ASD-like phenotypes.

**Figure 5.**
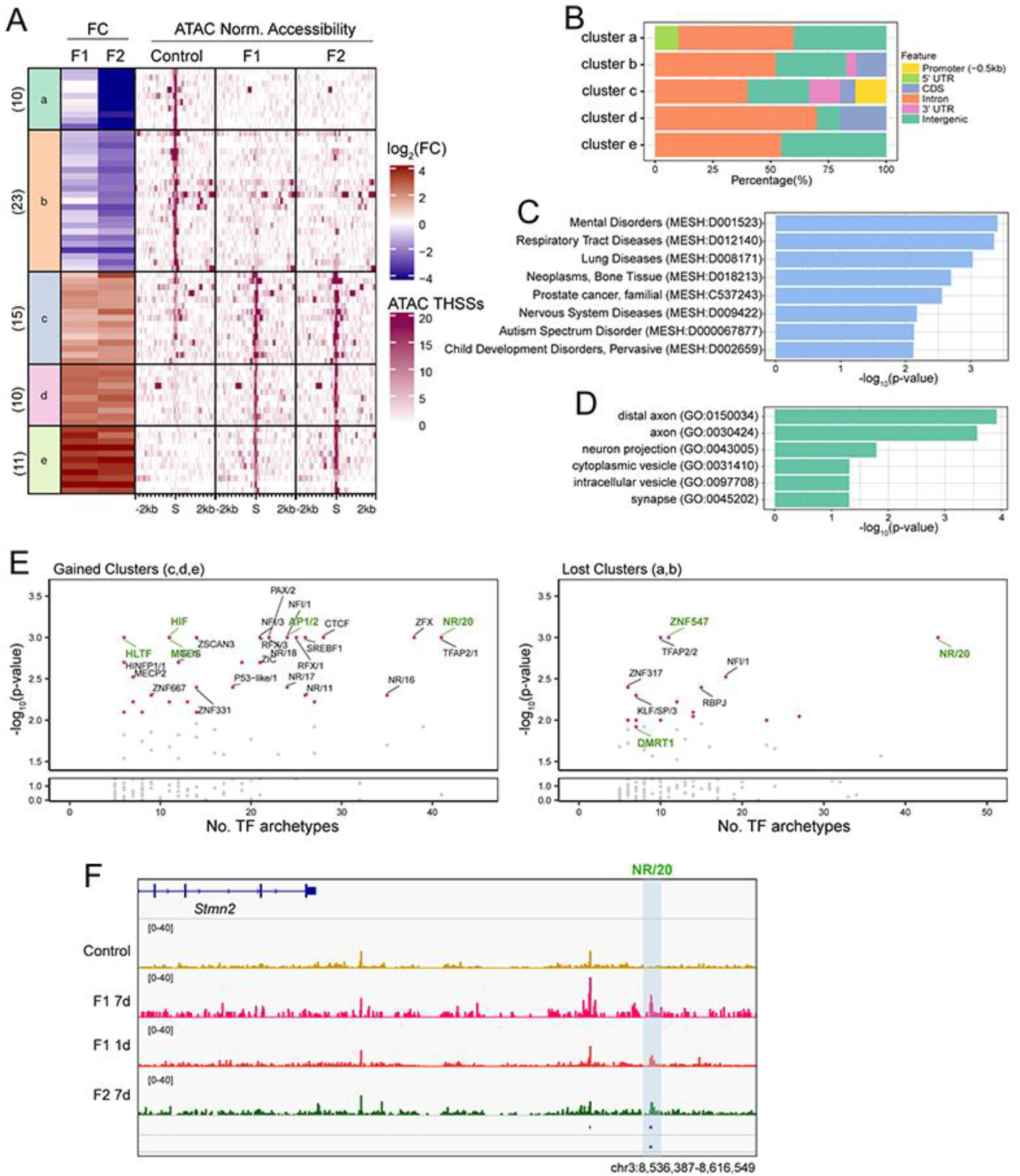
Sevoflurane induced alterations of TF binding maintained in sperm of F1 and F2 males. (A) K-means cluster analysis of THSSs (n=69) present in both F1 and F2 sperm but not control or vice versa. Fold change of pair-wise comparisons (shown on the left) of 7-day exposed F2 vs control was used in k-means clustering. The fold changes from the F1 sperm are shown on the left. The corresponding normalized subnucleosomal reads are shown on the right for control, 7-day exposed F1, and F2 datasets. In all clusters the alterations found in F1 sperm datasets are maintained in the F2 sperm. (B) Genome-wide distribution of THSSs present in the different clusters of panel A; promoters were defined as TSS −0.5 kb. (C) Enrichment analysis of disease ontology terms associated with the 69 differential THSSs shows genes involved in mental disorders, nervous system diseases, and ASD (Comparative Toxicogenomics Database, p-value <0.01, hypergeometric distribution and adjusted with Bonferroni correction). (D) Gene ontology of cellular component enrichment analysis associate the 69 THSSs with genes encoding proteins located at distal axons, cytoplasmic and intracellular vesicles, and synapses (PANTHER GO-SLIM, p-value < 0.01). (E) TF motif archetype enrichment analysis at differentially accessible clusters from panel A showing increased accessibility (clusters c, d and e; left) and lost accessibility (clusters a and b; right). TF motif archetype clusters with p-value less than 0.001 based on Monte Carlo permutation tests simulated genomewide were labeled with a pink dot. TF motif archetypes with p-value < 0.001 in the permutation tests simulated within the pool of ENCODE’s candidate cis-regulatory elements (cCREs) are marked in green. (F) Genome browser view of a 79 kb region upstream of the *Stmn2* gene. A differential THSSs with the NR/20 archetype is highlighted by a blue shaded box.

We then examined TF archetype enrichments at these 69 differential THSSs found in both F1 and F2 datasets (Figure 5F) and found some striking similarities with the F1 ASD dataset shown in Figure 3D. NR/20, which consist of Ar and GR binding motifs, is one of the most significantly enriched TF archetypes found in both gained and lost clusters and this association was also observed in the F1 ASD dataset (Figure 2D). This finding supports our hypothesis that Ar might be responsible for mediating the effects of sevoflurane on the germ cell epigenome. In addition to Ar, other nuclear hormone receptors present in NR/16,19,11,17,18 archetypes, including retinoic acid receptor gamma, estrogen and related receptors, are enriched in sites present in the gained but not lost clusters (Figure 5F, left). *Dmrt1* and *Znf547* TF archetypes are specifically enriched in the lost clusters. Dmrt1 is a gonad specific TF that is required for testicular development in vertebrates [76] and one of the genes frequently located in chromosome 9p deletions in humans [77, 78]. Most patients with 9p deletions share several common cognitive features, including intellectual disability, speech/language impairments, and ASD. Because the sevoflurane F2 ASD and F3 ASD males exhibit more severe impairments in social behavior compared to many published single-gene mutation animal models, the sperm of sevoflurane ASD animals may hold multiple clues to the origin of ASD pathology in the offspring. A gene found adjacent to differential THSSs present in both the F1 and F2 ASD datasets is *Stmn2,* which is highly expressed in the brain cortex (TPM > 100) (Figures 5G and S4). *Stmn2* encodes a member of the stathmin family of phosphoproteins that functions in microtubule dynamics and signal transduction, specifically in motor neuron growth and repair [79]. Changes in *Stmn2* expression have been associated with neurodegeneration, including Parkinson’s and Alzheimer’s disease (AD) [79-81]. Mutations in *Stmn2* lead to phenotypes similar to those found in ASD patients in human and animal studies, including anxiety behavior and impairment in social interactions [81-83]. Thus, our findings suggest that epimutations caused by exposing germ cells to sevoflurane can lead to ASD in the offspring, and these epimutations manifest as alterations in transcription factor binding near ASD genes that can be maintained in the male germline for multiple generations and transmitted both inter and trans-generationally.

## Discussion

The exponential increase in diagnosed cases of severe ASD in recent years demands a renewed effort on understanding the reasons for this surge. Emphasis has centered on using high throughput genomics to sequence the exome or whole genome of individuals with or without ASD symptoms and their families, and to try to identify sequence variants that correlate with ASD. This has led to the conclusion of an exclusively genetic basis for autism, with hundreds of coding mutations and thousands of non-coding variants suggested as the culprits for the development of ASD [1]. Often, these mutations are not present in the parents and arise de novo in the parental germline or in the somatic cells of the pre-implantation embryo. In parallel, it has been reported that exposure to dozens of chemicals present in the environment can increase the risk of ASD in both human and laboratory animals [1, 2, 5]. It is possible that these exposures lead to an increase in mutation rates or affect DNA repair processes. Alternatively, in addition to directly affecting brain development *in utero* or in early childhood, exposure to chemicals present in the environment may alter epigenetic information in the germline that could then affect gene expression in the embryo after fertilization. Observations reported here support this hypothesis and suggest that exposure of pregnant females to sevoflurane results in a dramatic alteration in the occupancy of specific TFs in the male germline at genes associated with ASD-like phenotypes. A subset of these altered sites is present in the sperm of directly exposed F1 animals and are maintained in the sperm of F2, correlating with ASD phenotypes in all three F1-F3 generations tested. Because in our experiments males were always outcrossed to unexposed control females, these observations suggest that the male gametes are sufficient to transmit epimutations initially induced by sevoflurane, both inter- and trans-generationally. It will be important to test whether exposure to sevoflurane can induce similar alterations in the female germline and whether ASD phenotypes can be transmitted to subsequent generations maternally through the oocyte.

The alteration in TF binding in the sperm of sevoflurane induced ASD males suggests that these altered sites could be epialleles responsible for the transmission of ASD phenotypes. However, it is important to note that not all exposed F1, F2 or F3 animals are autistic or have the same degree of severity of ASD phenotypes in response to sevoflurane exposure. This raises the possibility that sequence variants present in different individuals of the outbred CD-1 strain used in these studies determine the response of specific individuals in the population to sevoflurane exposure. The results suggest that sevoflurane exposure, and perhaps other environmental chemicals, act on the underlying genetic variability present in individuals of a population to elicit different disease outcomes.

Animal models have been used to knock out specific genes involved in autism in humans and recapitulate ASD pathology, including deletion or haploinsufficiency in *Shank3, Mecp2, Adpn*, and *Chd8* [84-89]. Interestingly, F1-F3 ASD mice arising from sevoflurane exposure crosses have more severe impairments in social interactions than single gene knockout mouse models of ASD. This may suggest that sevoflurane affects multiple pathways that contribute to ASD pathology compared to single gene ASD models. The mechanisms by which alterations in the distribution of TFs in sperm affect gene expression after fertilization are unclear. We have previously shown that occupancy patterns of TFs in sperm are similar to those observed in cells of the preimplantation embryo [40, 41], suggesting transmission of this information between generations. It is possible that these TFs affect gene expression when transcription begins in the 2-cell embryo, leading to the activation of other genes that initiate a transcriptional cascade and eventually lead to changes in gene expression in cell of the target organs, in this case the brain. Alternatively, altered occupancy of TFs may protect from DNA remethylation in the epiblast, and this unmethylated state serves as the epigenetic memory that allows novel patterns of gene expression in target organs later in development_[39, 67]. Our analyses focus on a specific window of embryonic development that may be very sensitive to sevoflurane exposure. During this developmental window, both the brain, which is undergoing neurogenesis, and the germline of the embryo that will give rise to the offspring, are exposed to sevoflurane. It is possible that postnatal or adult exposures have different effects on the brain and germline and act by mechanisms different from the ones proposed here. Although we mainly focused on assessing abnormalities in social and repetitive behaviors in sevoflurane-exposed mice, it would be interesting to examine if these animals exhibit locomotor abnormalities or AD-like cognitive-deficit phenotypes, since both ASD and AD have been connected through shared pathology, including memory deficits, cognition changes, demyelination, oxidative stress and inflammation [90], and GA exposure has been reported to have lasting impact on cognitive functions [14-23]. Given how widely anesthetic gases are utilized in surgical procedures, most often in critically ill pregnant women and pediatric patients, a full assessment in human subjects should be carried out to reduce the possible high risk of neurological impairments in the exposed subjects and subsequent generations.

## Acknowledgements

We would like to thank Dr. Cynthia Vied at the Translational Science Laboratory of Florida State University for help with Illumina sequencing, Dr. Yoon Hee Jung for help with the analysis of ATAC-seq data, and Dr. Michael Nichols for suggestions on the use of Monte Carlo permutation test analyses. Funding: This work was supported by the National Institute of Environmental Health Sciences of the National Institutes of Health under Award Numbers R01ES027859 and P30ES019776. HLW was supported by NIH F32ES031827. The content is solely the responsibility of the authors and does not necessarily represent the official views of the National Institutes of Health.

## Authors contributions

H-LW and VGC conceived and designed the study. H-LW planned and performed experiments, analyzed data, and contributed to writing the manuscript; SF performed experiments, analyzed data, and contributed to writing the manuscript. VGC planned experiments and contributed to writing the manuscript.

---

**Supplemental Figure S1.**
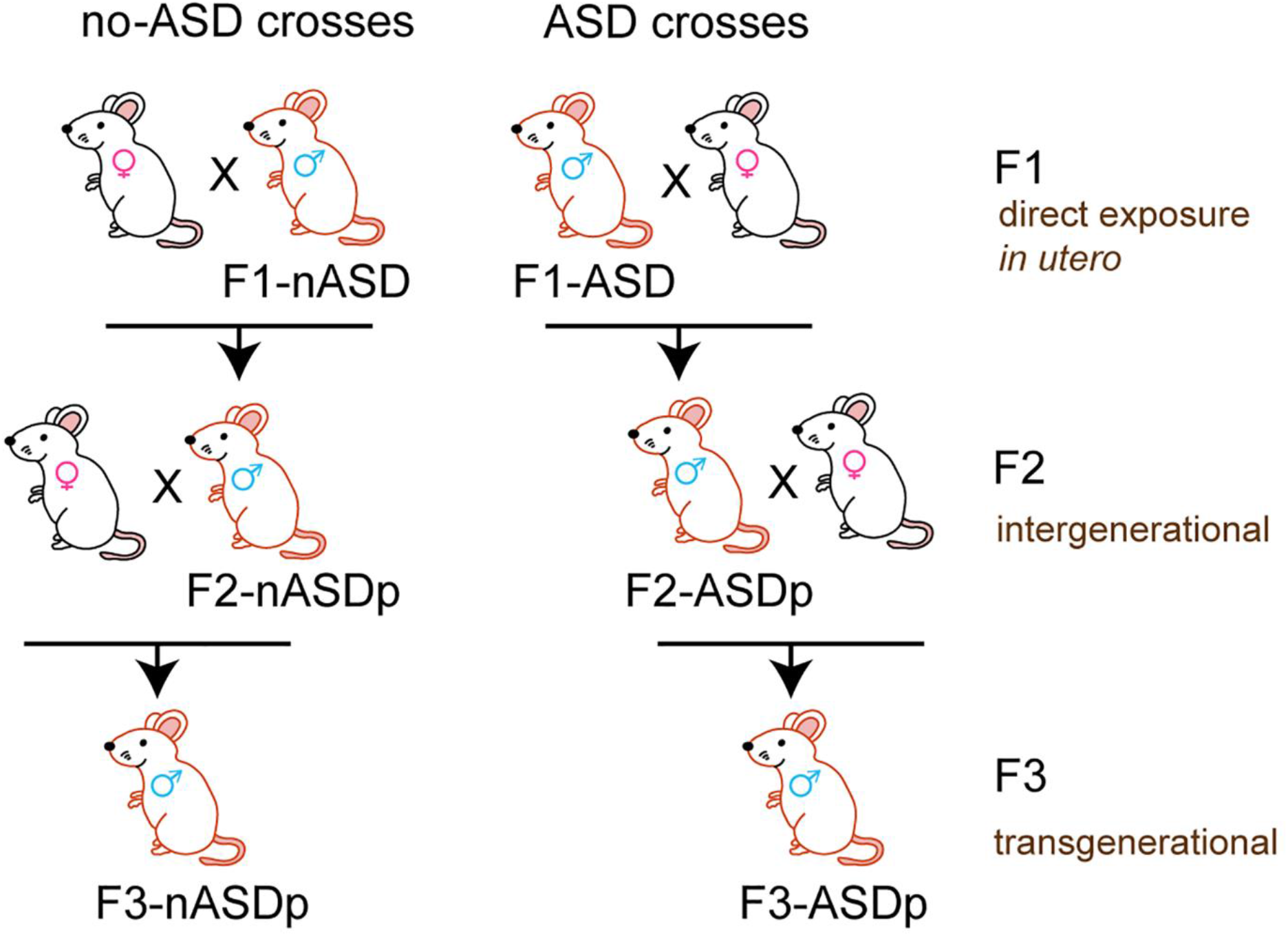
Breeding schemes of sevoflurane-exposed mice. F1 males directly exposed to sevoflurane *in utero* were crossed with control females to generate F2 animals. F2 is the inter-generational group, where the germline that gave rise to F2 was directly exposed to sevoflurane in the F1 embryo. Two separate crosses, each with two biological replicates, were performed to generate F2 animals that are sired by either F1 exposed male mice showing ASD phenotypes or showing normal behavior. Similarly, two separate crosses, each with two biological replicates, were performed to generate F3 animals that are either sired by no-ASD F2 or ASD F2 animals. F3 is the first trans-generational group that was never directly exposed to sevoflurane.

**Supplemental Figure S2.**
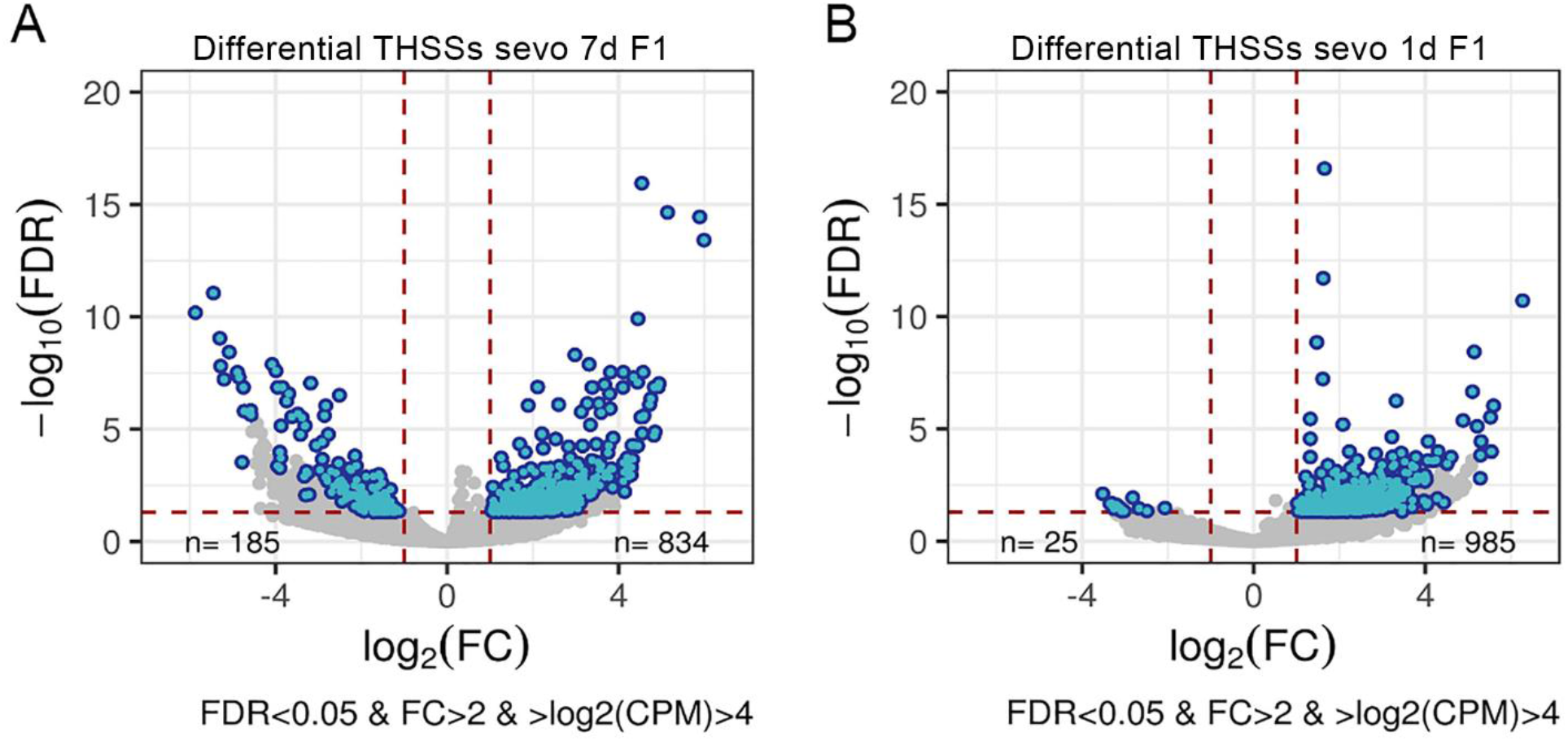
Analysis of differential THSSs between sevoflurane exposed F1 and control animals. Volcano plots showing pair-wised comparison between sperm THSSs datasets of 7-day exposed F1 (A) and 1-day exposed F1 (B) compared to control animals. The x-axis indicates the log_2_(fold change) and the y-axis shows the -log_10_(FDR). Differential THSSs with FDR<0.05, fold change (FC) >2 and log_2_(counts per million, CPM>4) are shown in blue.

**Supplemental Figure S3.**
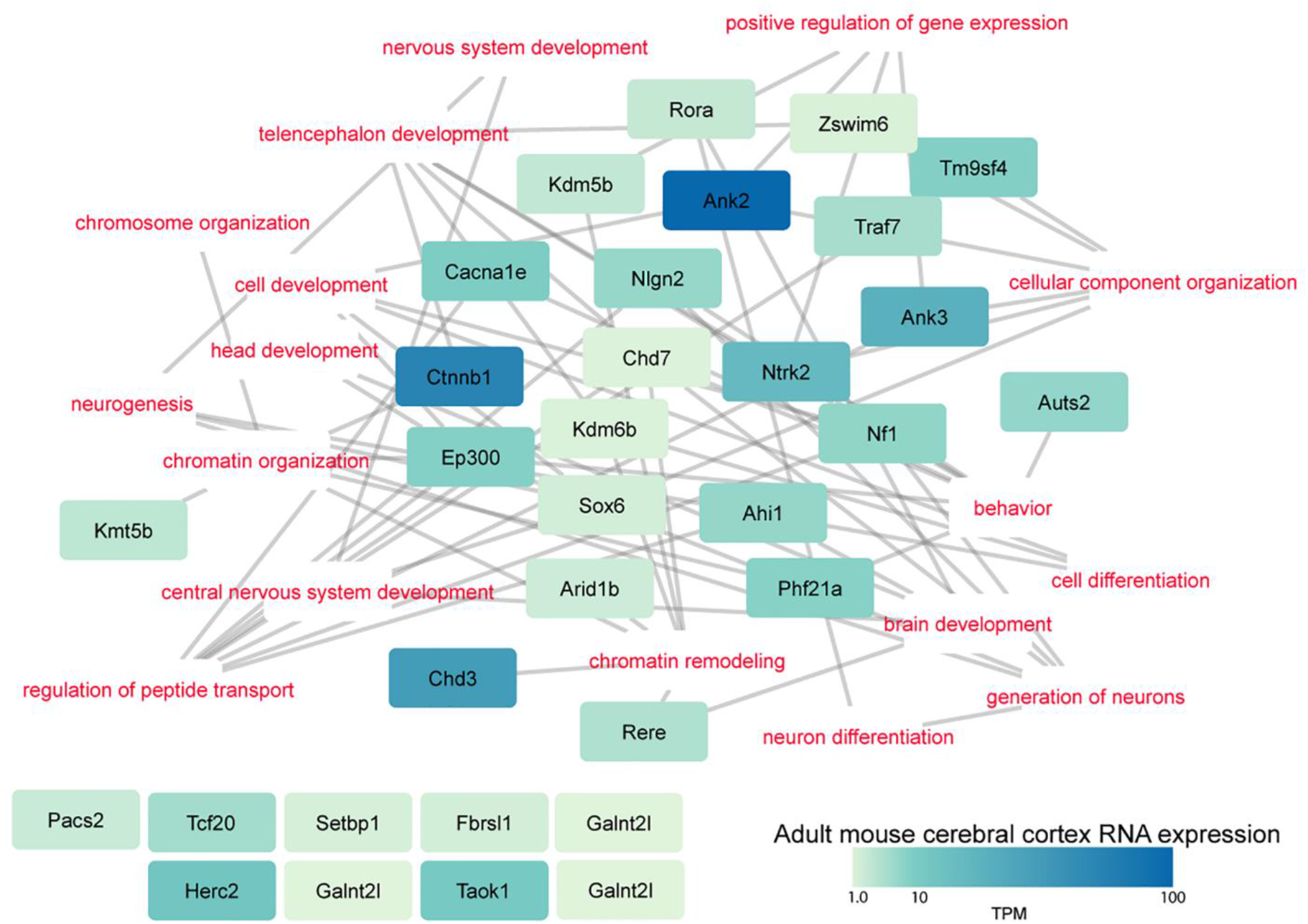
Genes implicated in ASD are enriched in/around differential THSSs between F1 exposed to sevoflurane and control. Gene ontology network analysis of genes adjacent to differential THSSs. Each node of the network contains genes previously shown to be mutated in ASD patients and enriched in/around THSSs differentially occupied in sperm of F1 sevoflurane exposed versus control males. Colors indicate levels of RNA expression in the mouse cortex in transcripts per million, TPM. All ASD genes are connected based on enrichment in gene ontology biological processes terms (marked in red). Genes with no shared ontology are indicated at the bottom left.

**Supplemental Figure S4.**
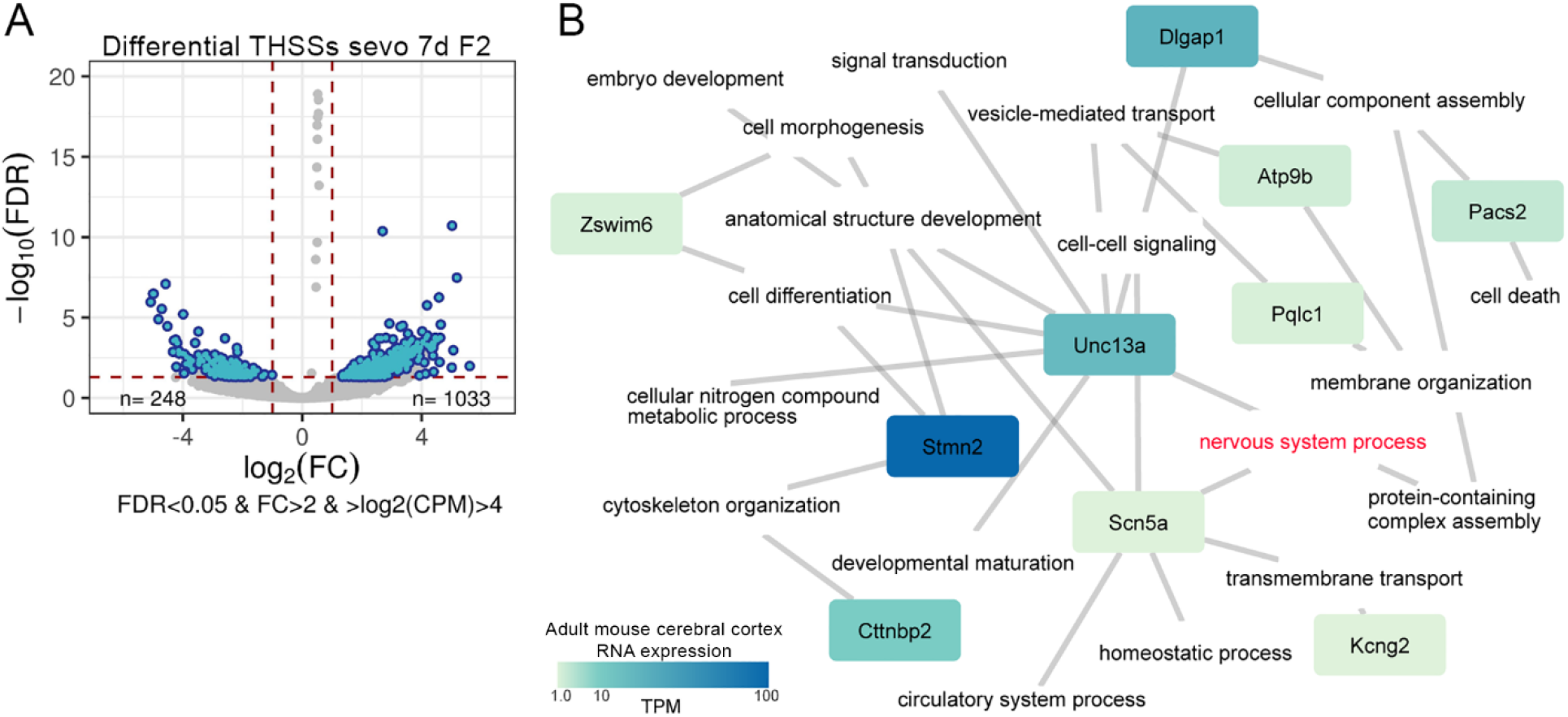
Genes involved in mental disorders and synapsis function are highly associated with differential THSSs shared between F1 and F2 mouse sperm. (A) Analysis of THSSs present in the sperm of F1 and F2 sevoflurane ASD males but not in control. Volcano plot showing pairwise comparison. The x-axis shows the log2(fold change) and the y-axis indicates the log10(FDR). Significant THSSs with FDR<0.05, fold change (FC) >2 and log2(CPM)>4 are shown in blue. (B) Mental disorders and synapsis function genes enriched in/around differential conserved THSSs are shown in each node of the network. Color intensity indicates RNA expression level in mouse brain cortex in transcript per million, TPM. All ASD genes are connected by gene ontology biological process terms. Nervous system processes term is marked in red.

